# Ecological Scaling of Temporal Fluctuations with Bacterial Abundance in Gut Microbiota Depends on Functional Properties of Individual Microbial Species and Bacterial Communities

**DOI:** 10.1101/2024.11.28.625948

**Authors:** Peng Su, Konstantine Tchourine, Dennis Vitkup

## Abstract

Macroecological relationships that describe various statistical associations between species’ abundances, their spatial, and temporal variability are among the most general laws in ecology and biology. One of the most commonly observed relationships is a power-law scaling between means and variances of temporal species abundances, known in ecology as Taylor’s law. Taylor’s law has been observed across many ecosystems, from diverse plant and animal ecosystems to complex microbial communities. While many mathematical models have been proposed to explain the potential origins of Taylor’s law, what determines its scaling exponents across species and ecosystems is not understood. Here, we use temporal trajectories of human and baboon gut microbiota to analyze the relationship between functional properties of individual bacterial species and microbial communities with the scaling of species-specific and community-level Taylor’s law. The species Taylor law characterizes – for each individual species – the relationship between the species’ temporal abundance means and temporal abundance variances across host organisms. On the other hand, community-level Taylor’s law characterizes – in each host organism – the scaling across multiple species between their temporal abundance means and temporal abundance variances. For community Taylor’s law, we find that the power law scaling is strongly associated with the microbiota abundance of certain nutrient-degrading enzymes in the gut. Notably, our results demonstrate that the availability of enzymes metabolizing starch glycogen significantly increases Taylor’s law scaling. We also find that species Taylor’s law depends on the individual species’ functional properties. Specifically, we observe lower Taylor’s law scaling for species with larger metabolic networks, for species that are able to grow on a larger number of carbon sources, and for species with particular metabolic functions, such as glutamine and folate metabolism. Overall, our study reveals that Taylor’s law scaling is strongly associated with the functional capabilities of bacterial communities and individual microbial species’ biosynthetic properties, which are likely related to their ecological roles in the gut microbiota.

## Introduction

Many recent studies have investigated the dynamic nature of the gut microbiota, revealing significant fluctuations in the abundance of bacterial species over time [1-3]. Notably, ecological dynamics of the gut play a critical role in shaping host health and disease, ranging from autoimmune diseases to colorectal cancer and to obesity-related diseases [4-7]. Understanding the biological mechanisms behind these temporal fluctuations is crucial, as they provide insights into the functional properties and the stability of the microbiota. Most recently, it was demonstrated [8-12] that it is possible to characterize the temporal variability of gut microbiota abundance using a set of quantitative macroecological scaling laws similar to those described previously in plant and animal ecologies [13-16]. One such relationship, Taylor’s law (TL), describes a power law association between the temporal variance *V* and the the temporal mean *M* of species abundance:

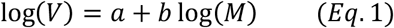

where *a* and *b* are the linear regression parameters of the power law fit [17-20]. In this study, we investigated the functional correlates of both the community Taylor’s law and the species’ Taylor’s law. The *community Taylor’s law* characterizes the scaling between species’ mean temporal abundances and species’ temporal abundance variances across species for each host organism, whereas the *species Taylor’s law* characterizes the scaling between species’ temporal abundance means and temporal abundance variances across host organisms for each species (Figure 1A). These scaling relationships were found in hundreds of insect and animal species [21, 22] and have been recently shown to be useful for identifying pathogenic and invasive species [11, 23] and understanding abundance dynamics in microbial ecosystems [24, 25]. Specifically, we recently showed that Taylor’s law can be used as a null model of temporal fluctuations, which enabled the identification of the species deviating from the law as likely biomarkers of disease and gut dysbiosis [11]. Taylor’s law has been used in ecology to measure both spatial and temporal dynamics, both at the species level and at the level of ecological community [11, 24]. It has also recently been observed in microecological systems such as human and mouse gut, human vaginal microbiota, and soil [11, 24, 25] with the slope *b* varying considerably across species and ecosystems.

**Figure 1.**
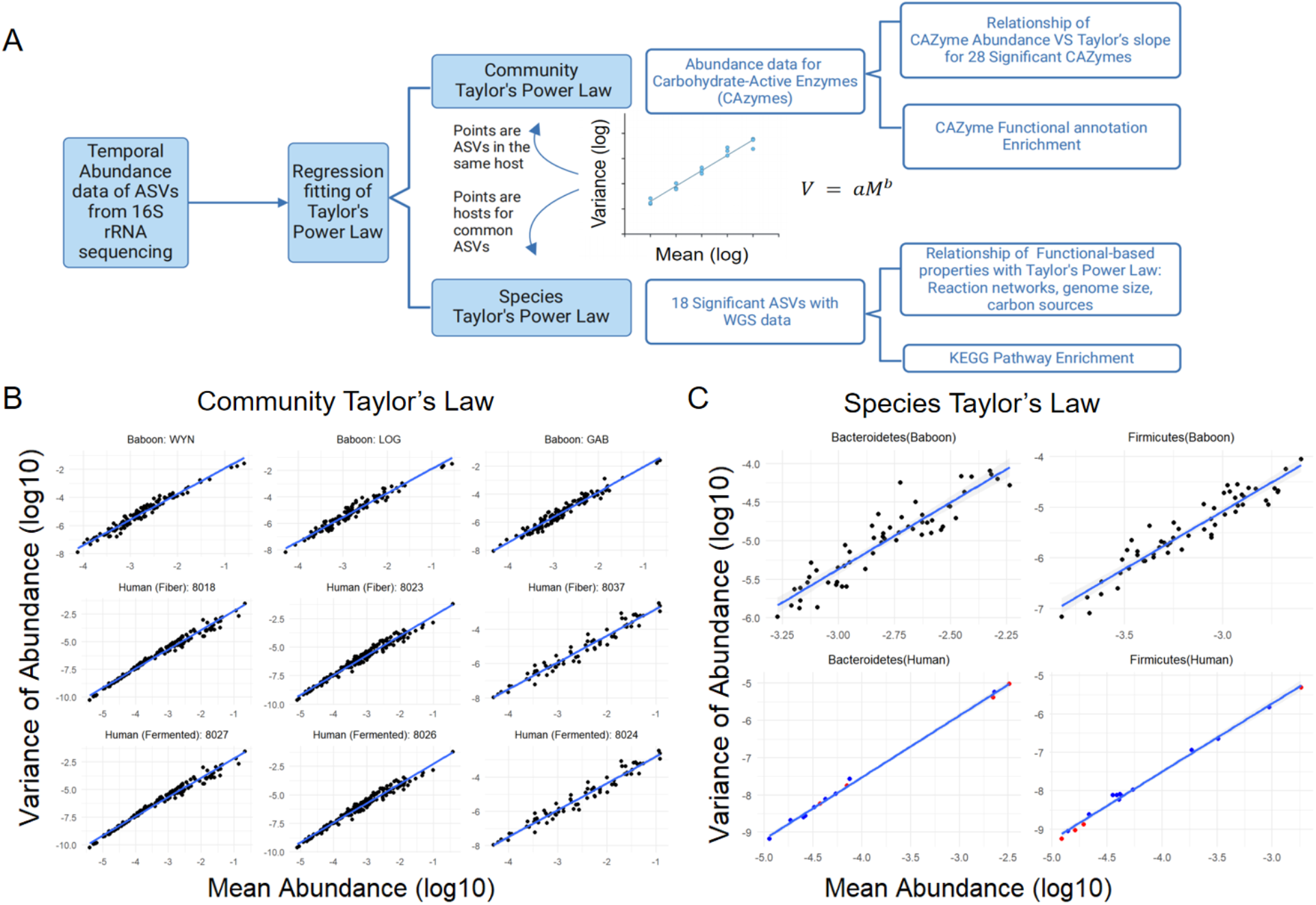
A. Outline of the analysis performed in this study. B. Representative community Taylor’s law fits in three baboon hosts and six human hosts fed a Fiber or Fermented food diet. C. Species Taylor’s law fits for representative Bacteroidetes and Firmicutes species in baboon and human data. The color indicates different diets for human samples.

Various theoretical explanations have been proposed to explain the variability of TL slope *b* [22, 26-28]. Some studies suggested that the expected slope for populations that fluctuate uniformly due to environmental variability is *b* = 2 [26, 29]. Notably, as we previously demonstrated for scaling values close to *b* = 1, species abundance changes in human and mouse gut microbiota are substantially larger in magnitude than the expected random fluctuations due to sampling noise [11]. Therefore, this scaling is not a result of sampling noise but represent an important characteristic of the underlying complex ecological dynamics. Experimental observations demonstrate that TL scaling coefficients are widely distributed across species and ecosystems [11, 19, 21, 24], and their mechanistic determinants and correlates remain unclear. A previous statistical analysis of time-series data has pointed to population-level characteristics, such as skewness, coefficient of variation, and synchrony, as correlates of the TL slope [30]. Additionally, computational simulations have proposed that biological factors like interspecies competition, resource distribution, and possibly environmental fluctuations can play a role in shaping abundance patterns over time [26-28]. However, biological correlates of Taylor’s law scaling, such as the functional properties of microbial communities and individual bacterial species, have not been empirically investigated.

In this study, we examined the association of biological properties of microbial species with the variability in Taylor’s law scaling in the context of the gut microbiota. Specifically, we analyzed a 10-year-long time series of baboon gut microbiota data [31], and a 14-week-long time series of human gut microbiota data [32]. The baboon dataset, collected in Amboseli National Park in Kenya, comprises over 5000 gut microbiota fecal samples, making it one of the longest microbiota time series to be analyzed using ecological scaling laws. The human dataset was collected on individuals following two diets (a plant-based fiber diet and a fermented foods diet) and includes metagenomic time-series measurements of carbohydrate-active enzymes (CAZymes), which code for enzymes that act on carbohydrate molecules [33], thus reflecting the overall carbohydrate processing capacity of the microbiota. We further mapped the Amplicon Sequence Variants (ASVs) in the human time-series dataset to known microbial genomes [34] (Figure 1A) and then utilized these mappings to investigate the relationship of Taylor’s law scaling with taxonomic composition, metabolic capabilities, and functional properties of bacterial communities and individual bacteria (Figure 1A). Collectively, our study demonstrates an important role of the functional properties of individual species and communities in determining the microbiota fluctuation scaling.

## Results

### Community Taylor’s Law Reflects Diet and Ecosystem-Level Functional Capabilities

In order to characterize the variability of Taylor’s law slopes across diverse gut microbial communities, we analyzed bacterial temporal abundance data from 56 baboon hosts measured on average over the period of eleven years and, separately, data from 36 human samples across nine time points spanning 14 weeks and two diets (Figure 1). We used the abundances of 16S rRNA Amplicon Sequence Variants (ASVs) to calculate abundance means and variances at the species level. We first focused on community TL, which characterizes the ecological dynamics of the entire bacterial community (Figure 1B, Figure S1). We found that for baboons, typical Taylor’s law slopes ranged from *b* = 1.7 to *b* = 1.9 (Figure 2A), with a mean of 1.811 ± 0.005; we observed no significant difference between male and female baboon community TL (Kolmogorov-Smirnoff test, *p* > 0.1). Community TL slope distribution across the 36 human hosts overlapped with the distribution in baboons but was significantly lower (Wilcoxon rank-sum *p* <1 x 10^-14^), ranging from *b* = 1.4 to *b* = 1.8 (Figure 2B). We also observed that human subjects consuming a fiber-based diet had a slightly but significantly (Wilcoxon rank-sum *p* < 0.05) lower mean community TL slope (*b* = 1.57 ± 0.03) than the subjects on a fermented food diet (*b* = 1.62 ± 0.02) (Figure 2B). Notably, the difference in TL scaling distributions between human and baboon datasets corresponds to a lower variance among high-abundance species in human data compared to baboons (Figure S2A). The power law scaling of the average abundance ranks of ASVs with their average abundances (“rank-abundance scaling”) was also more evenly distributed in baboons than in humans (Figure S2B). This suggests that the differences in community TL distributions between the two datasets are not merely due to different collection time intervals (typically 1.5 weeks for humans and 1.5 months for baboons) but may also reflect the diverse patterns in microbial community structures between the baboon and human host species.

**Figure 2.**
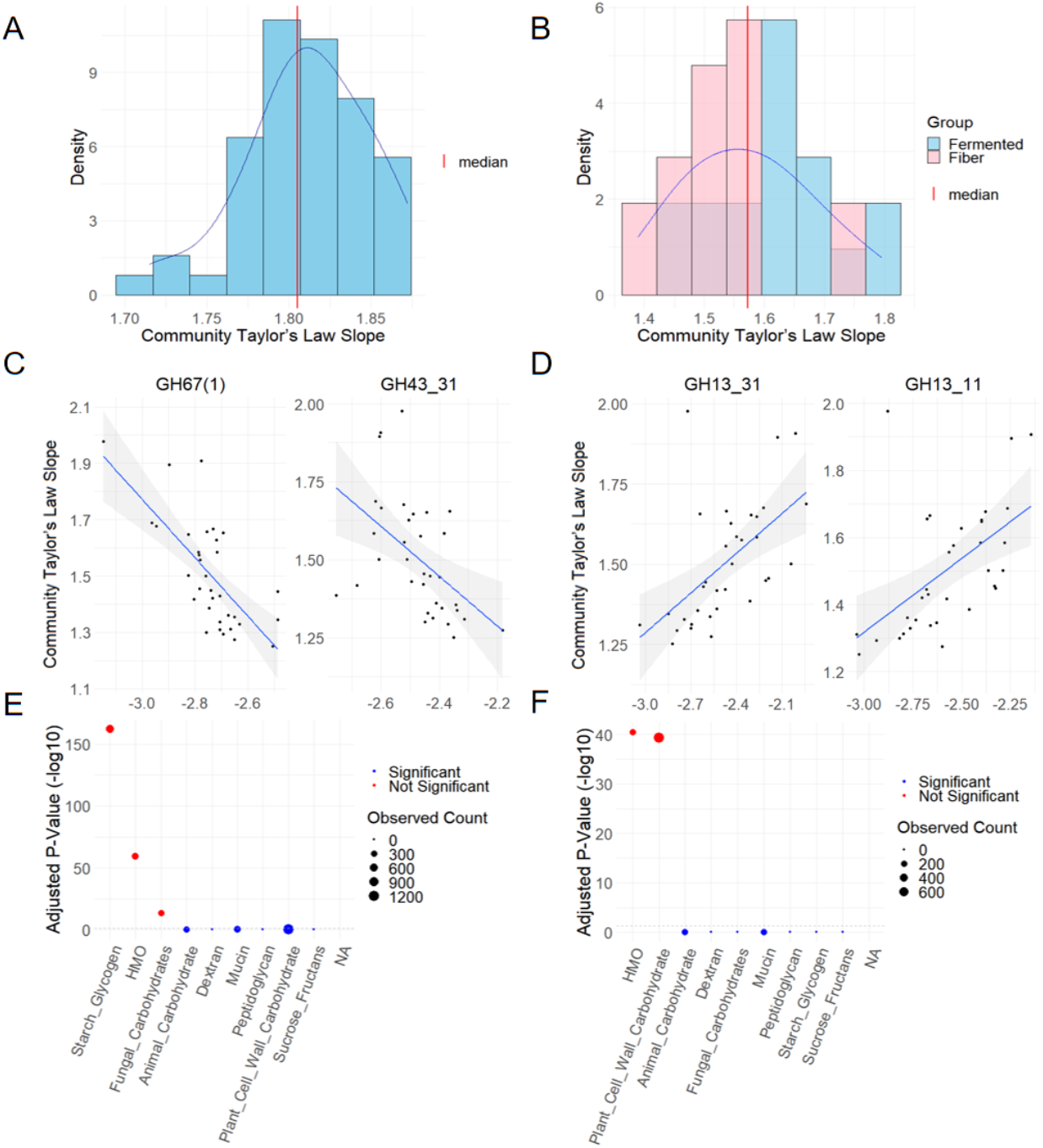
A-B. The distribution of community Taylor’s law slope for (A) 56 baboon hosts and (B) 36 human hosts. Different colors in (B) indicate different diets of human hosts. Vertical lines represent median values for each group. C-D. The relationship between average CAZyme abundance and community Taylor’s law slope across human hosts for two example CAZymes annotated as (C) Starch Glycogen Degradation and (D) Plant Cell Wall Carbohydrate Degradation. E-F. Significant CAZyme categories enriched among CAZymes with positive (E) and negative (F) correlations with community Taylor’s law slope. Shaded areas in C-D represent the 95% confidence interval of the linear fits.

Differences in host diet can lead to changes in metabolite availability in the gut [35, 36]. Previous metagenomics analysis of the human gut microbiota found that fiber and fermented diets affect the abundance of Carbohydrate-Active Enzymes (CAZymes) [32]. To determine how the functional properties of the microbiota may mediate the effect of diet on community TL in humans, we quantified the capability of the microbial community to digest individual carbohydrates using the measured temporal profiles of Carbohydrate-Active Enzyme (CAZyme) categories, averaged across the metagenomics time series for each human subject. To that end, for each of the 654 CAZymes with non-zero abundances in at least 18 humans, we calculated the correlation between the CAZyme abundance and the community TL slope across human hosts. We found that 28 CAZymes showed a significant correlation with the community TL slope (Benjamini-Hochberg corrected *p* < 0.05 ), with 20 CAZymes exhibiting positive correlations and 8 CAZymes exhibiting negative correlations (Figure 2C-D). Positively correlated CAZymes were enriched in starch glycogen degradation (Figure 2E, hypergeometric test with Benjamini-Hochberg corrected *p* < 0.05 ), suggesting that higher community TL slopes correlate with an increased ability of the bacterial community to digest starch-rich foods, such as grains and cereals [37]. In contrast, CAZymes that had a significant negative correlation with community TL slopes were enriched in plant cell wall carbohydrate degradation (Figure 2F, hypergeometric test with Benjamini-Hochberg corrected *p* < 0.05 ), with seven of the eight negatively correlated CAZymes belonging to this category. Overall, these results demonstrate that, although community Taylor’s law scaling is not substantially influenced by the host diet, the ability of the host’s gut bacterial community to break down and synthesize particular metabolites does have a strong and significant effect on community Taylor’s law scaling.

### The Broad Distribution of Species Taylor’s Law Scaling Depends on Functional Properties of Individual Microbial Species

In contrast to community TL, which reflects properties of the entire ecological community within each host, species TL characterizes how temporal fluctuations of individual bacterial species scale with the abundance of that species across hosts (Figure 1A). For both the human and the baboon datasets, we used ASVs as a proxy for bacterial species abundances to calculate species TL for each ASV (Figure S3). We found that in baboons, species TL slopes ranged from 0.5 to 2.6 (Figure 3A), with a mean value of 1.67 ± 0.04, with all 126 ASVs exhibiting a significant power law relationship between temporal abundance means and temporal abundance variances (BH-corrected Pearson correlation test *p* < 0.05). Similarly, across human subjects, species TL slopes ranged from 0.5 to 2.1 (Figure 3B), with the mean slope of 1.53 ± 0.02, and with all 215 ASVs also showing a significant power law relationship between abundance means and variances (BH-corrected Pearson correlation test *p* < 0.05). We first asked if the wide variability in species TL scaling is due to some species demonstrating relatively noisier TL scaling fits. To that end, we calculated the coefficient of determination of the fitted Taylor’s law, i.e. the fraction of the variance of ASV abundance variances explained by the mean abundance. Both in baboon and human datasets, we found that indeed, ASVs with weaker scaling (lower R-squared) tend to have lower TL slopes (Figure S4), suggesting that lower values of species TL slopes are at least partially explained by lower-quality power law fits. However, we note that for baboon data, TL fits with a relatively high-quality scaling of *R*^2^ > 0.6, making up 92% of baboon ASVs, had TL slopes ranging from 0.9 to 2.1. Even more strikingly, in human data, even the highest-quality TL fits (R^2^ > 0.9), which made up 75% of all human ASVs, had a broad distribution of TL slopes, ranging from 1.1 to 2.1. Thus, the variability in species TL slopes across ASVs is not merely a consequence of the quality of the Taylor’s law fits but likely represents real biological differences in the scaling between species.

**Figure 3.**
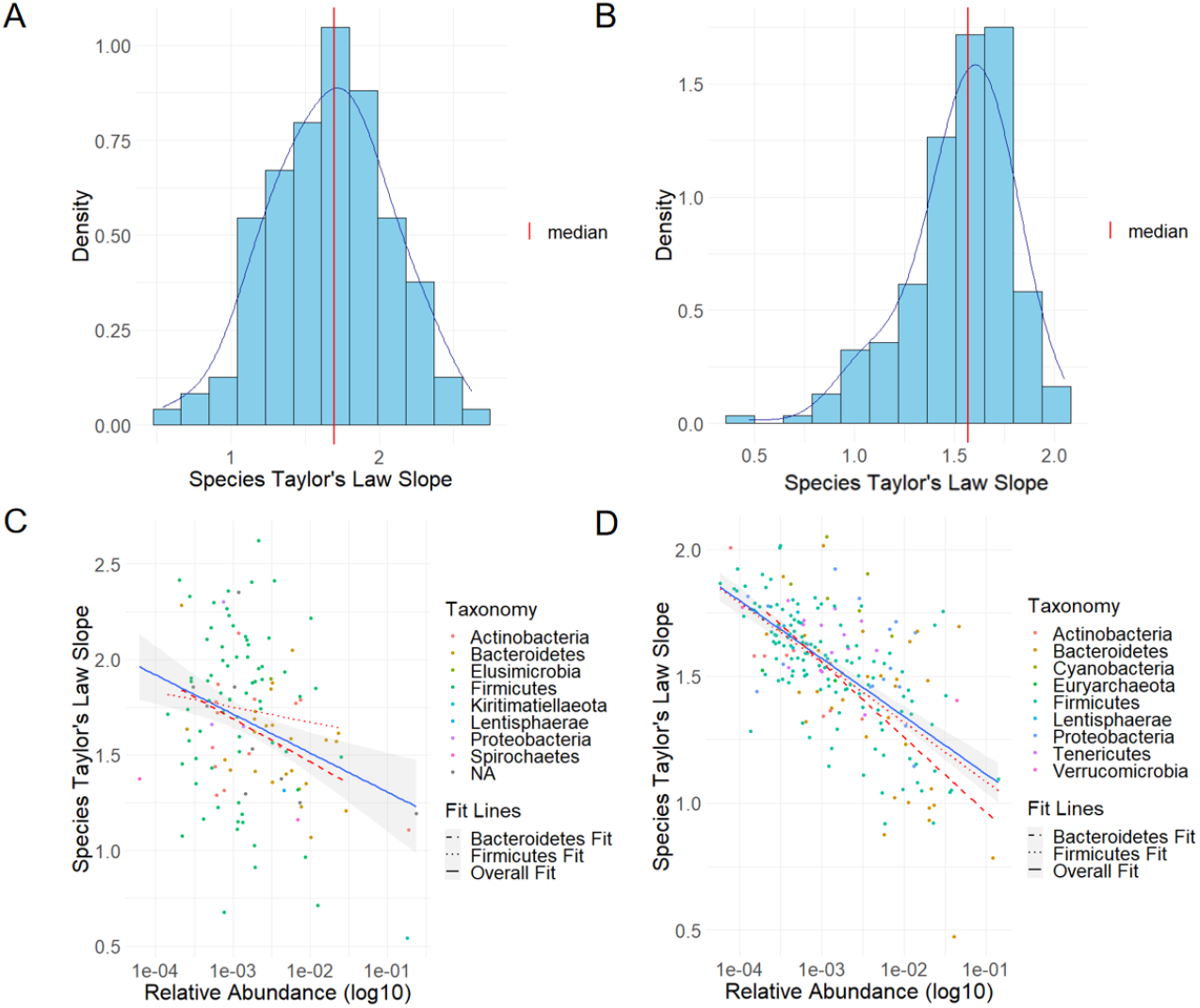
A-B. The distribution of species Taylor’s law slopes for (A) the 126 baboon ASVs and (B) the 215 ASV in the human data. C-D. The relationship between relative abundances of ASVs and their species Taylor’s law slopes for (C) the baboon data and (D) the human data. The solid blue lines in C-D indicate the linear fit based on data points from all species, whereas the red lines represent fits to Firmicutes (dotted lines) and Bacteroidetes (dashed lines).

We next asked whether species TL is indeed a property of individual microbial species or if the scaling depends primarily on the collection of considered host subjects. To that end, we assessed the robustness of species TL slopes to the selection of host subjects by randomly separating the 56 baboon hosts into two equal groups and comparing the species Taylor’s law scaling across the same species, calculated within each independent group of hosts. We repeated this procedure for 100 random groupings of host subjects and found a strong and significant correlation of TL slopes between the two random groups for all ASVs (mean Pearson’s *r* = 0.709 ± 0.005, all BH-corrected *p* < 0.05) (Figure S5A). Analogously, randomly splitting the 36 human subjects into two equal groups and comparing the species TL slopes calculated across 100 random samplings resulted in a substantial and significant correlation between the two groups of independent human hosts for all human ASVs (mean Pearson r = 0.499 ± 0.008, all BH-corrected p < 0.05) (Figure S5B). Notably, splitting the human hosts by diet (18 hosts in Fiber diet and 18 in Fermented) resulted in a TL slope correlation (Figure S6, Pearson r=0.5, *p* < 10^-5^) highly similar to the one calculated based on two randomly sampled groups of human host subjects independent of their diets. This suggests that the host’s diet does not play a significant role in determining species TL. The downsampling of baboon subjects from 56 to 36 hosts to match the number of subjects in humans resulted in similar correlations between random groups of subjects (Figure S5C), indicating that lower correlations in humans may be a result of a lower number of available subjects. In summary, these results suggest that species TL scaling is largely a property of the species themselves and possibly their functional characteristics, rather than the host, and that species TL scaling does not depend strongly on host diet.

### Species Phylogeny Does Not Explain Substantial Variation in Taylor’s Law Scaling

In order to investigate how various ASV properties affect species TL scaling, we first assessed the role of species taxonomy in determining species TL slope. We started by comparing average species TL slope differences at different levels of shared taxonomic categories (Figure 4A-B). We did not find a strong relationship between TL slope differences and the number of shared taxonomic categories, both for baboons (Spearman *ρ* = 0.01, *p* > 0.05) and humans (Spearman *ρ* = −0.02, *p* < 0.003), and average TL slope differences between different hierarchical categories were small compared to the variance of TL slopes within each category. Similarly, the correlation between 16S sequence differences and TL slope differences between ASVs was small (Spearman *ρ* = −0.04, *p* < 2 × 10^-5^ for baboons and *ρ* = 0.02, *p* < 0.001 for humans).

**Figure 4.**
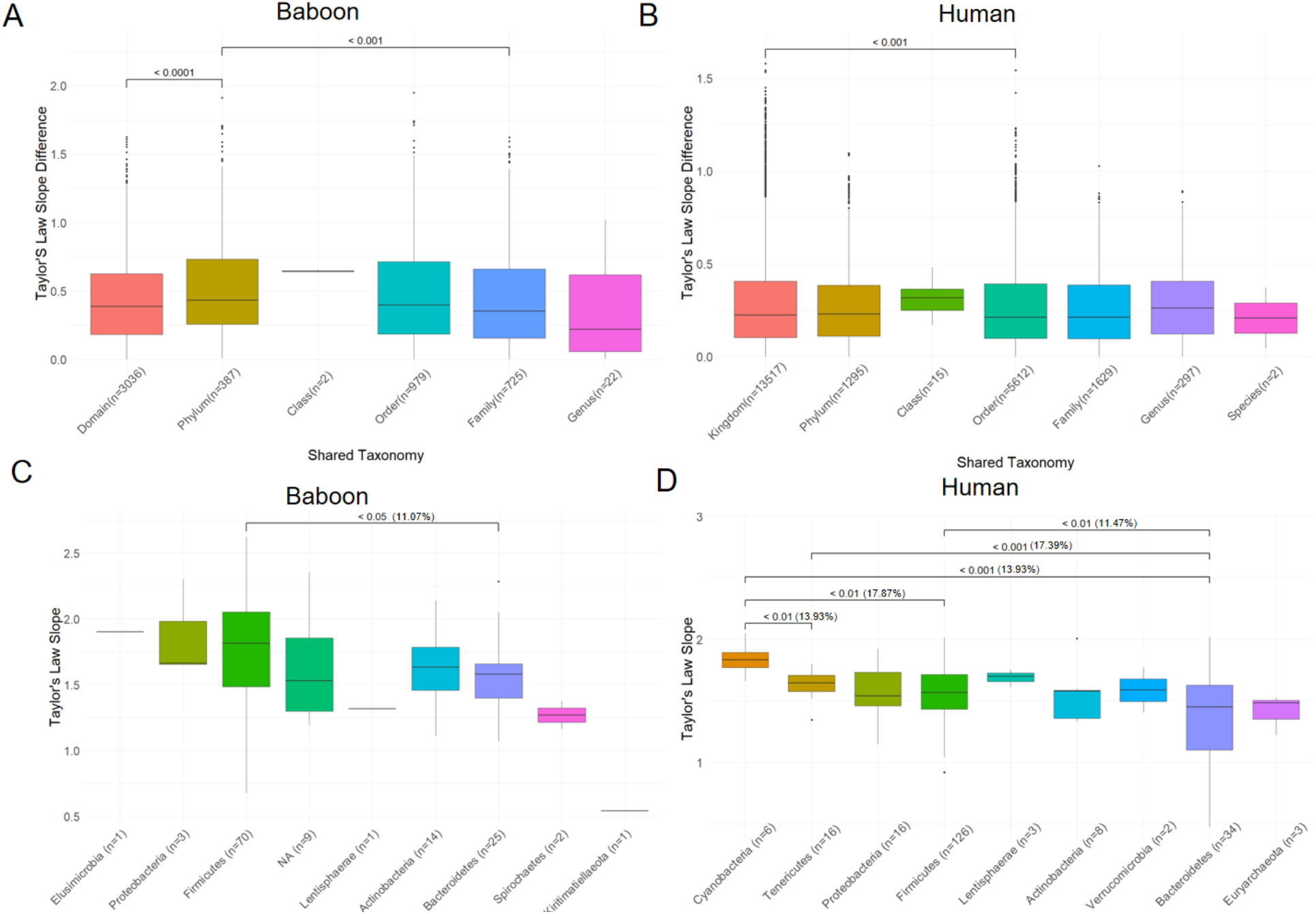
A-B. The relationship between the phylogenetic distance between pairs of ASVs and the difference in species Taylor’s law slopes between the corresponding ASV pairs. Each box blot indicates the lowest taxonomic level shared in the baboon (A) or the human data (B). C-D. The distribution of species Taylor’s law slopes in different phyla for (A) the baboon data and (B) the human data. All significant Wilcoxon test p-values for pairs of phyla are shown (followed by percentage difference in mean).

We next performed a more detailed assessment of the role of phylogeny in determining species TL scaling by separating all ASVs based on phylum and comparing species TL law distributions across the phyla (Figure 4C-D). In both baboon and human datasets, we found that Bacteroidetes have significantly lower TL slopes (mean TL slope 1.37 ± 0.06 in humans and 1.56 ± 0.05 in baboons) than Firmicutes (mean TL slope 1.54 ± 0.02 in humans and 1.75 ± 0.05 in baboons; Wilcoxon test *p* = 0.014 and *p* < 0.01, respectively). In humans, Cyanobacteria ASVs had the highest species TL slopes, ranging from 1.7 to 2.1, which was significantly higher than both Firmicutes (Wilcoxon *p* < 0.002 ) and Bacteroidetes (Wilcoxon *p* < 0.0003 ). Considering that Firmicutes and Bacteroidetes are known to have substantial differences in their average abundances [11], we next tested the dependence of species TL slopes on the abundance of ASVs within and across phyla. We found that higher abundance baboon ASVs tend to have lower TL slopes across Bacteroidetes (Pearson r=-0.46, *p* < 0.05), across all ASVs (Pearson r=-0.31, *p* < 10^-3^), but not across Firmicutes (Figure 3C). Similarly, in humans, higher abundance ASVs were found to have lower TL slopes overall (Pearson r=-0.65, *p* < 10^-5^), as well as across Bacteroidetes (Pearson r=-0.61, *p* < 0.05) and Firmicutes (Pearson r=-0.70, *p* < 10^-16^) (Figure 3D). Moreover, the close similarity of the linear fits between abundances and species TL slopes for Firmicutes and Bacteroidetes suggests that the average differences in TL slope distributions between these two phyla in humans are likely due to the differences in their abundance distributions.

### Properties of Species’ Metabolic Networks Strongly Influence Taylor’s Law Scaling

To further investigate how the functional and genomic properties of bacteria, such as metabolic network features and the ability to grow on various nutrients, affect their individual species TL scaling, we next mapped 16S rRNA sequences from each ASV to whole genomes of bacterial isolates from the human gut available in the CAMII biobank [34]. Filtering the matches at >97% similarity resulted in 18 human ASVs that can be reliably mapped to high-quality whole genomes of human gut bacterial species. Using these mappings, we found that across these 18 ASVs, genome size strongly correlated with species TL scaling (Pearson r=-0.48, *p* < 0.05), with larger genomes corresponding to significantly smaller TL slopes (Figure 5A). To test whether this relationship may reflect the effect of bacterial metabolism on the ecological dynamics of each species, we next used modelSEED, a popular tool for the reconstruction, analysis, and simulation of metabolic networks [38], to generate the metabolic reaction network for each ASV (see Methods). The investigation of the modelSEED networks for the 18 bacteria showed that the sizes of metabolic networks had an even stronger correlation with species TL slopes than did their genome sizes (Figure 5B, Pearson r=-0.61, *p* < 0.005), suggesting that the number of metabolic reactions available to a bacterial species may have a strong effect on the dynamic ecological properties of the species. Notably, the relationship between the size of the metabolic network and the TL scaling was mostly independent of the species abundance (partial Pearson correlation r=0.5, *p* < 0.04), whereas the relationship with genome size was no longer significant after controlling for bacterial abundance. This demonstrates that reaction network size, and thus likely the overall metabolic complexity of each bacterial species, has a strong influence on species TL scaling.

**Figure 5.**
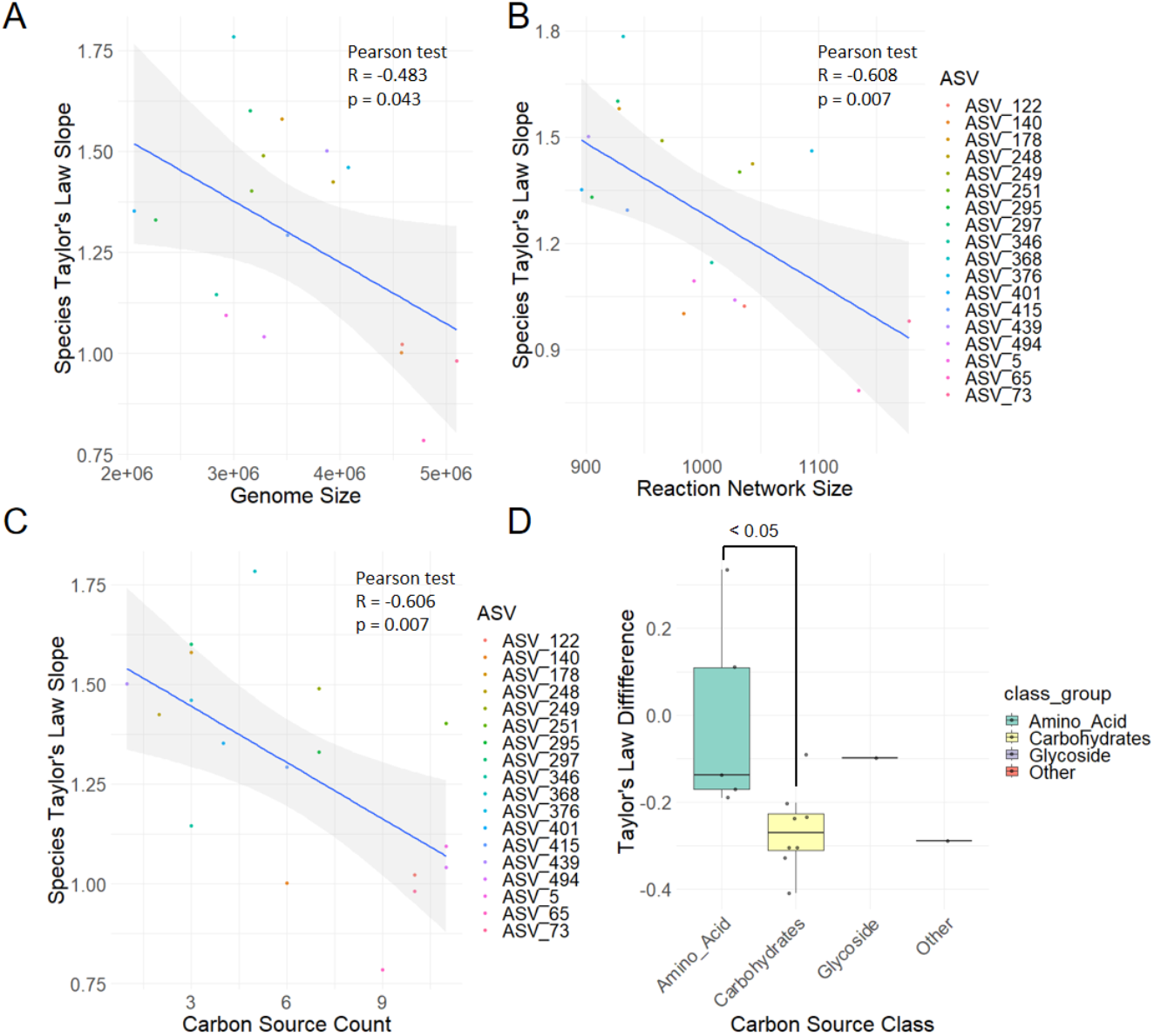
A-C. The correlation of species Taylor’s law slope with (A) genome size, (B) reaction size, and (C) the number of distinct carbon sources that can support the growth of each ASV. Shaded areas represent the 95% confidence interval of the linear fits. D. Distributions of the difference in Taylor’s law slope between ASVs that can and cannot grow on a given carbon source (points) for different carbon source categories.

To test whether the effect of metabolic network size on species TL scaling is related to the ability of each ASV to synthesize biomass from various metabolites, we next used Flux Balance Analysis (FBA) and the constructed modelSEED models [39] to test the ability of each of the 18 ASVs to grow on 24 distinct and commonly used carbon sources (see Methods). Interestingly, we found that ASVs capable of utilizing a broader range of carbon sources for growth tend to exhibit significantly lower TL slopes (Figure 5C, Pearson r=-0.61, *p* < 0.01). This effect was also mostly independent of the ASV abundance (Pearson partial correlation between carbon source count and species TL slope, while controlling for ASV abundance, r=-0.50, *p* < 0.0 5). Moreover, this effect was also largely independent of the metabolic network size (Pearson partial correlation between carbon source count and species TL slope, while controlling for metabolic network size, r=-0.47, *p* < 0.05), suggesting that it is the metabolic capacity of the bacterial species, rather than simply the size of the reaction network, that primarily influences ecological Taylor’s law scaling. When tested the ability to grow on various carbohydrate and amino acid sources, we found that the ability of ASVs to grow on carbohydrate carbon sources was more strongly associated with a lower species TL slope than the ability to grow on an amino acid (Figure 5D, Wilcox *p* < 0.02).

### Species Taylor’s Law is Associated with the Functional Properties of Each Species

Because of the strong dependence of species TL slopes on metabolic network size and the number of carbon sources that can be utilized for growth, we further investigated which reactions and metabolic pathways are most strongly associated with species TL scaling. To that end, we considered ASVs with and without each metabolic reaction. We then compared the distributions of species TL slopes between the two groups. This analysis revealed 114 metabolic reactions that were significantly associated with TL scaling (Benjamini-Hochberg adjusted Wilcoxon *p* < 0.05), with effect sizes, i.e., the percent difference of TL scaling when a reaction was either present or absent, ranging from 24% to 39%.

To explore the biological role of reactions associated with species TL slope differences, we mapped all reactions in each ASV’s genome to known KEGG metabolic pathways [40-42]. Specifically, for each ASV, we determined how many reactions from each KEGG pathway are present in the reconstructed reaction network of the ASV. We found that the completeness of nine KEGG pathways was strongly correlated with species TL slope across ASVs after correcting for species’ abundance (Figure 6, Figure S7, Benjamini-Hochberg adjusted Pearson correlation *p* < 0.1 ). Notably, the completeness of folate cycle, sphingolipid metabolism, and alanine, aspartate, and glutamate metabolism pathways, corresponded to lower TL slopes (Figure S7 B-C). This result is consistent with the hypothesis that generalist bacteria with more complete networks, and thus higher carbon source flexibility, tend to have lower TL slopes. The only KEGG category which had a positive correlation between pathway completeness and species TL slope was related to the metabolism of Anthocyanin (Figure S7A), a compound present in certain plants [43], indicating that bacteria with specialized metabolic roles may have higher TL scaling. Overall, these results suggest that the variability in bacterial species Taylor’s law scaling significantly correlates with the differences in the metabolic capabilities of these bacteria.

**Figure 6.**
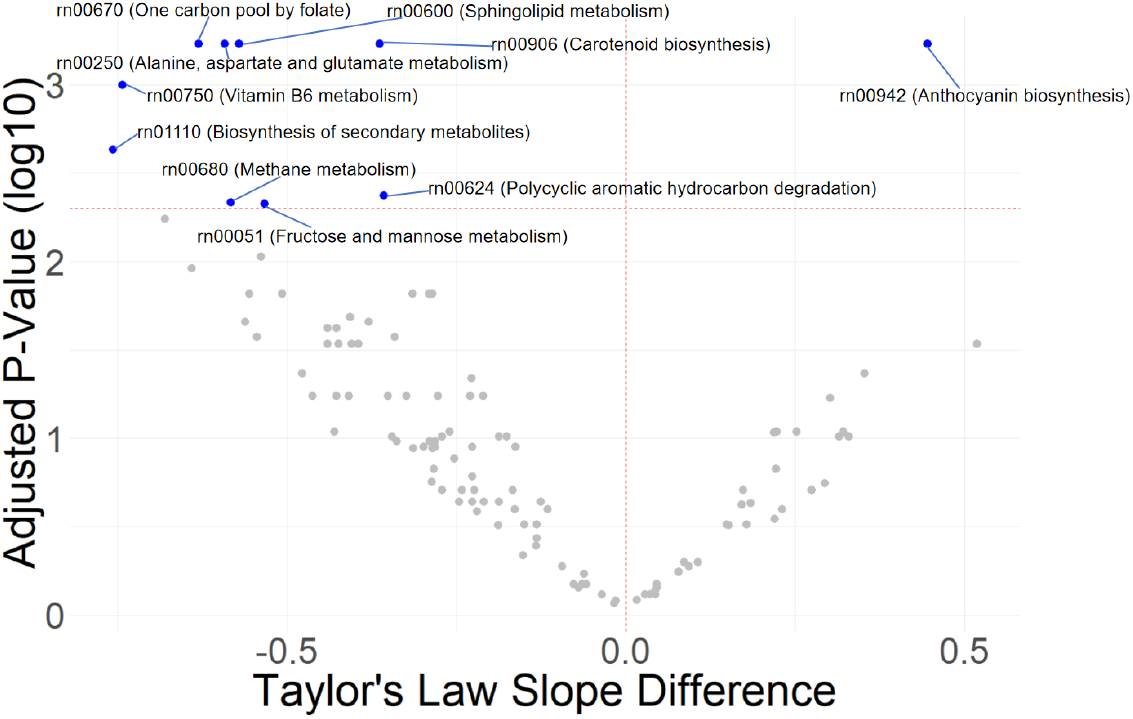
Taylor’s law slope differences between KEGG pathways. For each KEGG pathway (points), the p-value of the linear fit between pathway completeness and Taylor’s law slopes across species is plotted as a function of the total difference in Taylor’s law slopes between species with maximum and minimum completeness of that pathway, calculated using the linear fit (see Figure S7). Significant pathways are shown in blue.

## Discussion

In this study, we showed that the variability of Taylor’s law scaling – both across microbiota in different hosts and across species in the same host – depends on the functional metabolic properties of these microbial ecosystems and individual bacterial species. Certain functional categories, such as CAZymes involved in starch glycogen metabolism, can explain up to 43% of the variability in community Taylor’s law slopes, in contrast to host gender or the host diet, which were not strongly associated with community Taylor’s law scaling. Similarly, specific functional categories explained 25-39% of the variation in species Taylor’s law scaling, a higher fraction of the variance than was explained by species taxonomy.

Interestingly, we found that functional categories that have strong negative correlations with species Taylor’s law scaling tend to be involved in central carbon metabolism. We also observed that ASVs with larger metabolic networks and ASVs that can utilize a large number of carbon sources for growth have substantially lower Taylor’s law slopes, even after correcting for ASV abundances. This may be due to the differences in interspecies interaction patterns: bacteria that can utilize a large number of different carbon sources are less likely to engage in competition for resources compared to highly specialized bacteria. Indeed, species interactions through competition have been previously linked to Taylor’s law slope in simulations of microbiota dynamics [26-28].

In the broader context of ecology, we hope that our analysis will serve as an important step towards connecting macroecological scaling laws to functional and metabolic properties of individual species as well as functional properties of entire microbial communities. It would also be interesting to investigate in the future how functional and metabolic properties of bacteria influence other macroecological relationships, such as long-term drift [11, 13, 44, 45], short-term fluctuations [11, 13, 14, 46, 47], and residence and return times [11, 13, 48, 49]. In combination with emerging computational models of ecological communities [28, 34, 50-52] and the analysis of synthetic bacterial communities [53, 54], exploring how functional properties of individual species affect their macroecological relationships will be necessary to understand the ecological dynamics of living systems.

## Methods

### Data download and processing

ASV temporal abundance data from 56 baboon hosts measured over an average of eleven years [31] and data from 36 human samples across nine time points [32] was downloaded from the supplementary data of the corresponding studies. To account for technical noise, for the baboon dataset we only included ASVs having non-zero abundance in at least 20% of samples for each host, which resulted in 126 common ASVs for calculations of both community and species TL. For calculating community and species TL in the human dataset, we filtered ASVs to those having non-zero abundance in at least 20% of time points within each host and that are present in at least 10 different hosts, which resulted in 215 ASVs for the human dataset.

Taxonomic categories and 16S sequences for each ASV were downloaded along with their abundance profiles, as reported in Roche *et al*. and Wastyk *et al*. [31, 32]. To determine the correlation between phylogeny and Taylor’s law slope, ASV pairs with shared taxonomic categories were defined as ASVs with identical (non-NA) taxonomy, and 16S alignment differences were evaluated by the Hamming distance, where pairwise Hamming distances for each sequence pair were computed by counting the number of mismatched positions, excluding positions with missing data.

CAZyme abundances with functional annotations in the human dataset were also provided by Wastyk *et al*., 2022 [32], with CAZyme abundance time points corresponding to the ASV abundance time points. To evaluate the influence of relative CAZyme abundance on community TL, we selected data from 34 human subjects with at least two measurements for each CAZyme. For each selected host, only the CAZymes with non-zero abundance in more than half of the time points were included in the analysis. After identifying 28 significant CAZymes with strong correlations between average abundance and their community TL slopes, enrichment analysis was performed using the hypergeometric test to evaluate the probability of observing specific annotations among significant CAZymes, given their frequency among annotations for all 654 CAZymes. For each annotation, we assessed whether it was overrepresented among significant CAZymes relative to its occurrence in the total CAZyme pool. P-values were calculated for each annotation and subsequently adjusted for multiple testing using the Benjamini-Hochberg procedure at 10% false discovery rate.

### Isolates mapping

Human ASVs were mapped to human gut microbial isolate genomes from the CAMII database [34] at a 97% similarity level. For ASVs that mapped to multiple isolates, we restricted matches to high-quality isolates with at least 80% completeness, less than 5% contamination, and purity over 75%. We then picked the isolate with the highest completeness, lowest contamination, and highest purity, in that order, as the unique representative for that ASV. In total, 18 ASVs were successfully mapped to unique isolates, and their whole genome sequencing data was further used for metabolic modeling.

### Bacterial reaction networks and functional assignments

Metabolic models were built for the selected 18 ASVs using modelSEED [38], and biological functions and reactions were collected from these models for each ASV. For each reaction, we separated the 18 ASV models into two groups, based on whether or not this reaction is included in each model. We then compared the species TL distributions between the two groups of ASV models and used the Wilcoxon unpaired test to determine whether these distributions are significantly different. For the analysis of KEGG pathways, we used KEGG pathway annotations assigned to reactions by modelSEED. For each ASV model, the number of reactions that mapped to each KEGG pathway was compared to the species TL slope of that ASV. We used the partial Pearson correlation test, controlling for average ASV abundance, to determine which KEGG pathways are significantly correlated with species TL slope across the ASV models. For each KEGG pathway, we used linear regression to assign a linear model to the relationship between its pathway completeness and species TL slope, and the total effect size of that KEGG pathway was then calculated as the difference in its linear model’s prediction of TL slope between the ASV with the highest pathway completeness and the ASV with the lowest pathway completeness.

### Modeling of growth on carbon sources

We used the COBRA toolbox in MATLAB [55] to calculate FBA models for the 18 ASVs models, using as their only source of extracellular carbon each of the 24 distinct carbon sources (CSs) for which experimental data was available in Plata *et al*. [56]. Other carbon compounds were limited to a maximum combined uptake of 10 mmol of carbon per gram of dry weight. For each tested CS, transport and exchange reactions were added to each ASV model, and the ability of each ASV to utilize these CSs was assessed by optimizing the FBA model, with biomass flux serving as the metabolic objective for bacterial growth. Upon balancing the fluxes for each CS, biomass fluxes were normalized to a scale from 0 to 100, corresponding to the minimum and maximum flux values, respectively, across all CSs in each ASV model. Based on GENIII Biolog Phenotype Microarrays data for 40 diverse microbial species across 24 growth conditions [56], we observed that the normalized biomass fluxes of these 40 species exhibited a bimodal distribution. Building on this observation, we applied a Gaussian Mixture Model (GMM) and used the Bayesian Information Criterion (BIC) to determine the optimal cutoff, 37% of maximal growth, for this bimodal distribution, which was subsequently used as the threshold, to classify whether an ASV could grow on the provided carbon sources (CSs). Under these conditions, an ASV was considered capable of growing on a specific CS if its normalized biomass exceeded this threshold.

## Acknowledgements

This work was supported by the grants by from the National Institutes of Health (grant numbers R35GM131884 and R01DK118044).

## Supplementary Figures

**Figure S1.**
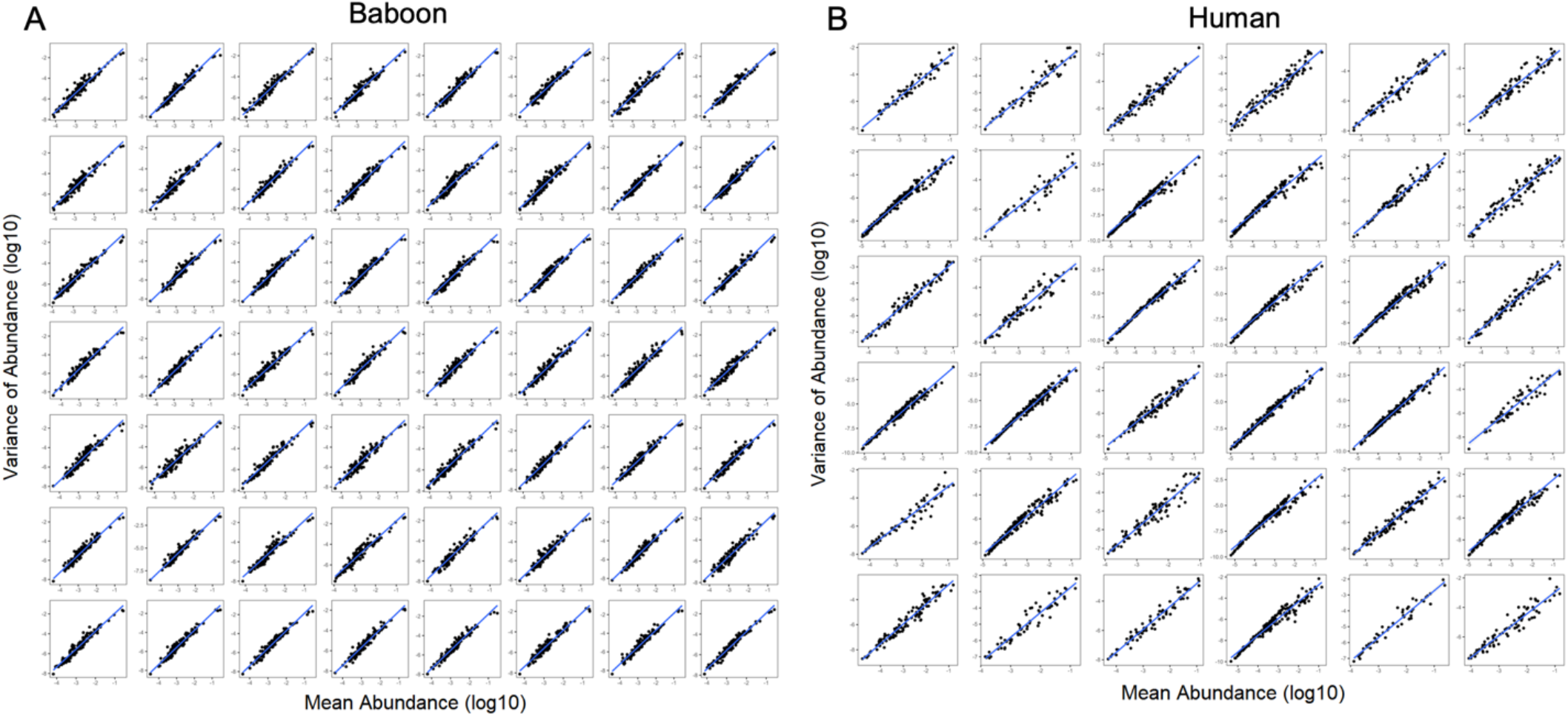
Community Taylor’s law fits for (A) all 56 Baboon hosts and (B) all 36 Human hosts.

**Figure S2.**
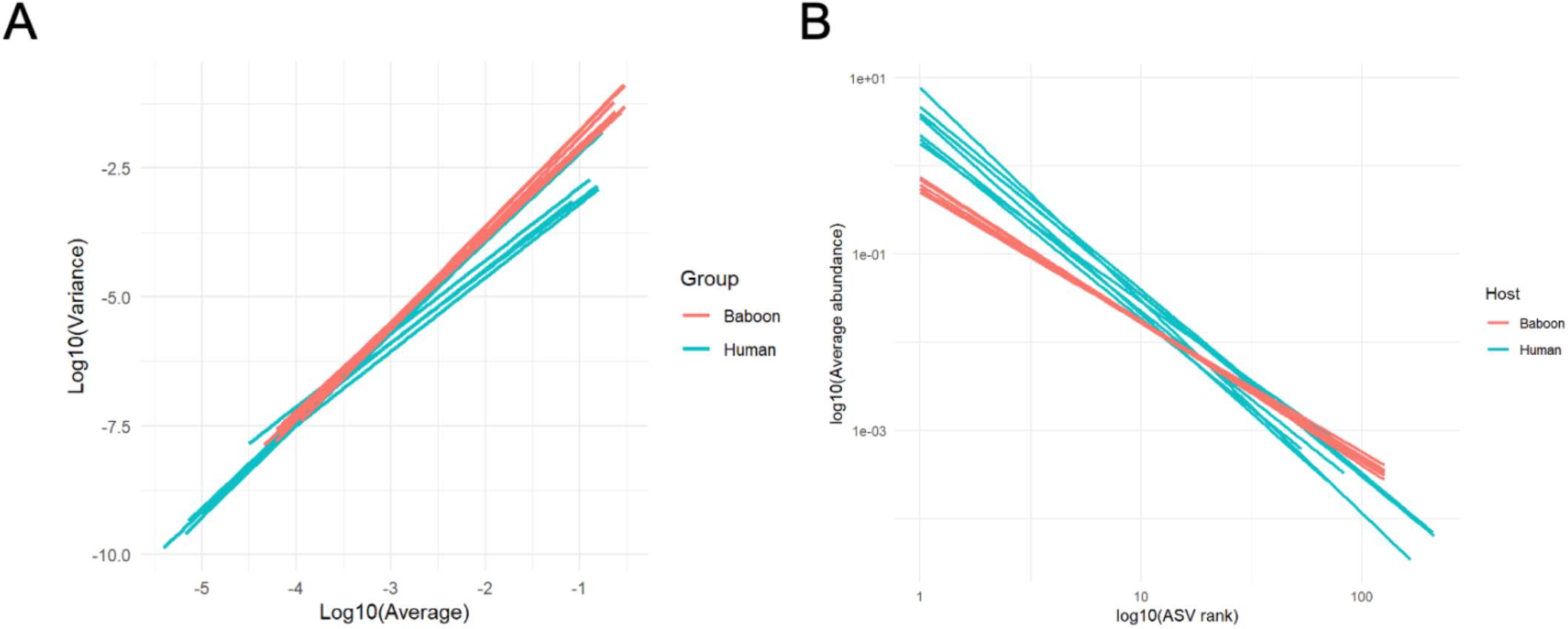
A. Community Taylor’s law fits for all baboon (red) and human (blue) hosts. B. Rank-abundance scaling fits for all baboon (red) and human (blue) hosts.

**Figure S3.**
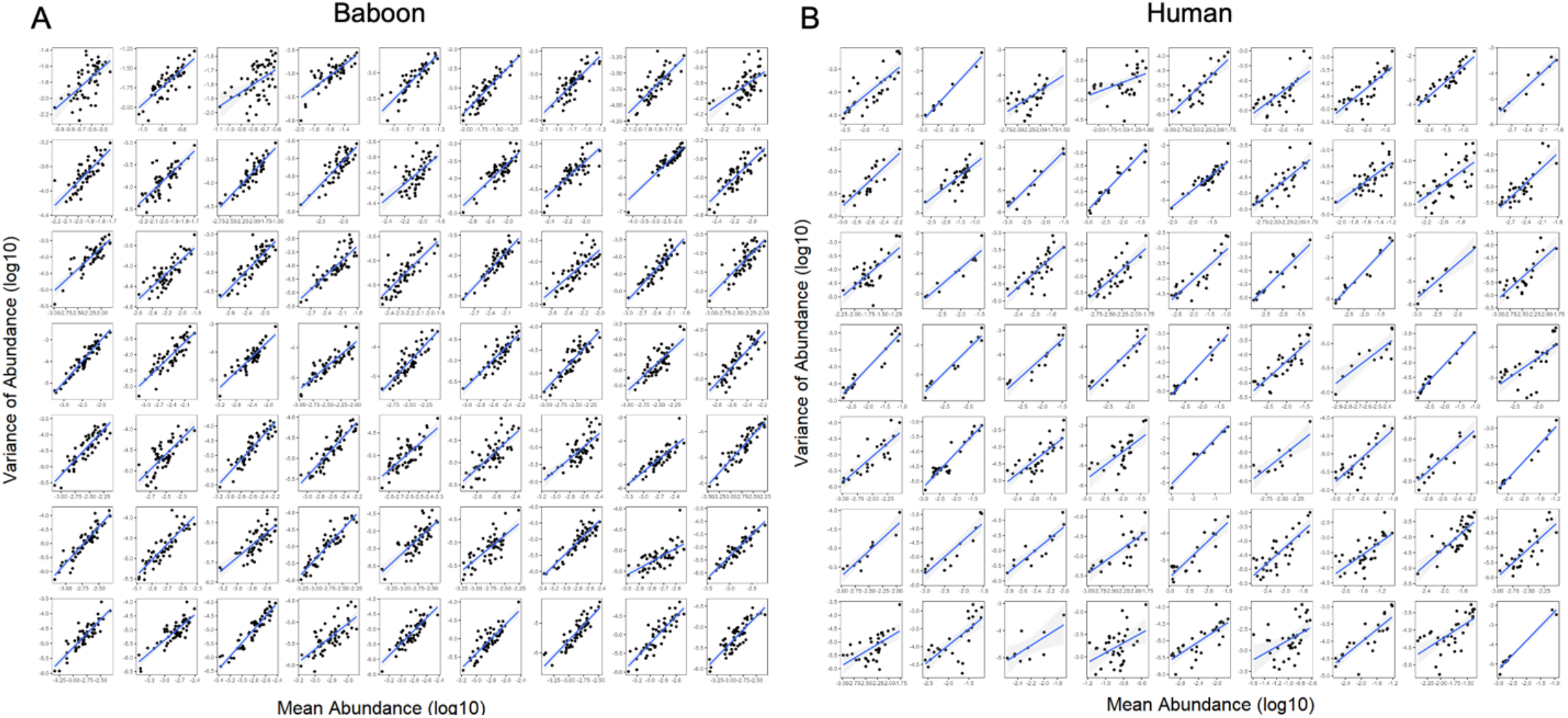
Species Taylor’s law fits for (A) 64 ASVs with the highest average abundances across all baboon hosts and (B) 64 ASVs with the highest average abundances across human hosts in which they were present.

**Figure S4.**
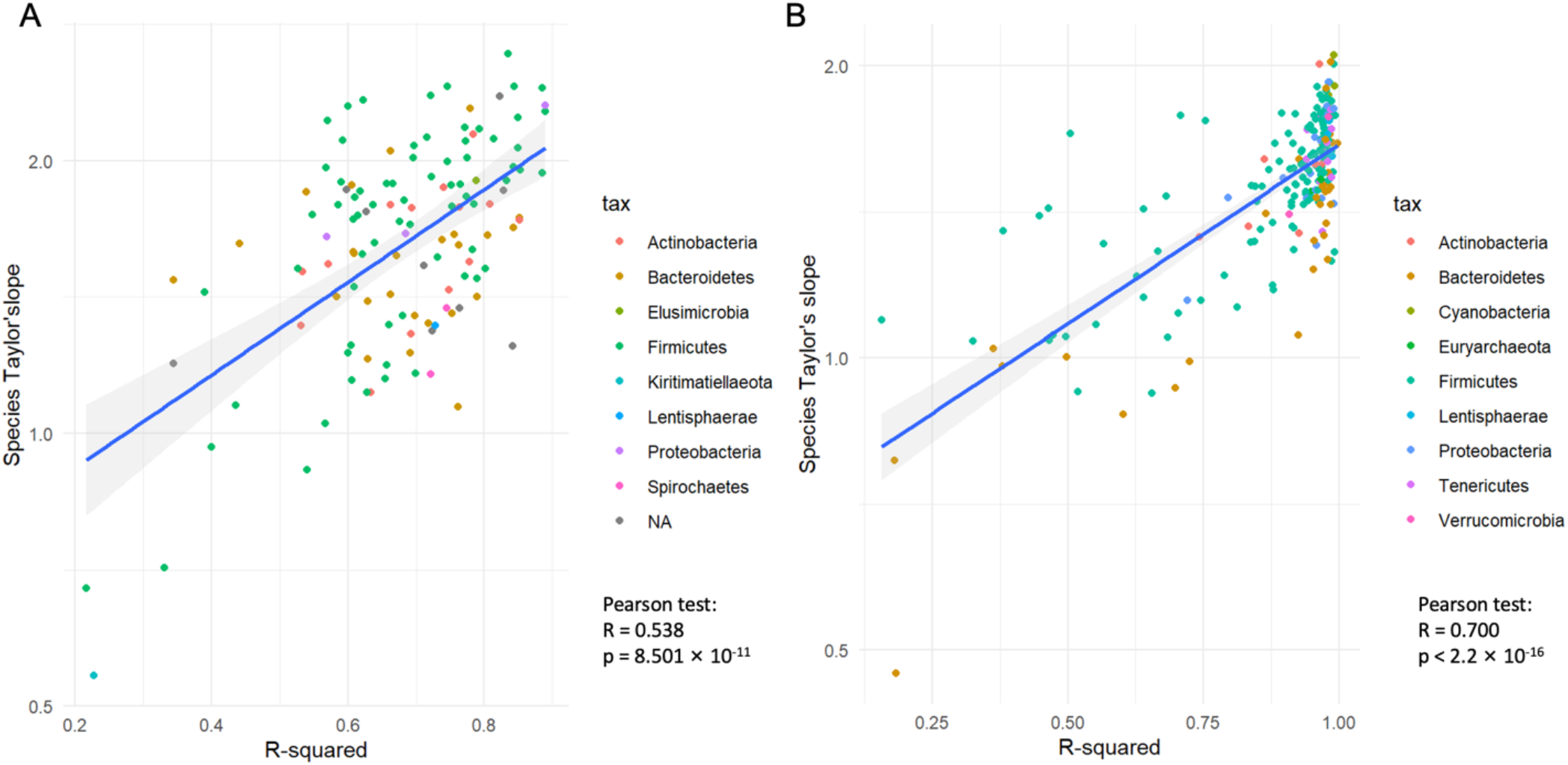
The relationship between individual Taylor’s law slope and the R-squared of the fitted linear correlation of the species Taylor’s law for (A) all Baboon ASVs and (B) all Human ASVs. Each point represents an ASV, and the color correspond to its phylum.

**Figure S5.**
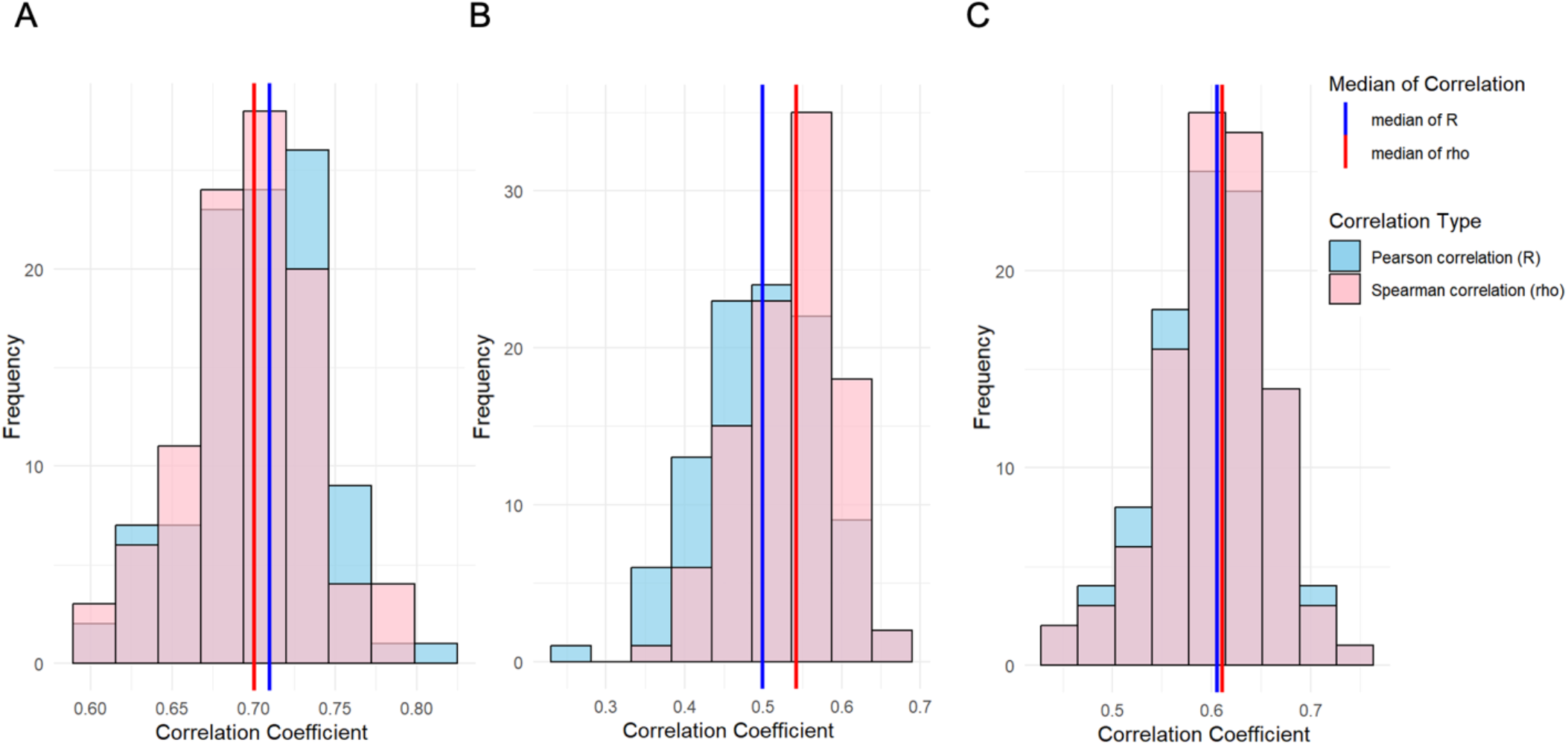
The distribution of correlations of species Taylor’s law slopes between two equal randomly-generated sets of hosts, across 100 random splits of (A) the Baboon data, (B) the Human data, and (C) the Baboon data downsampled to 36 hosts.

**Figure S6.**
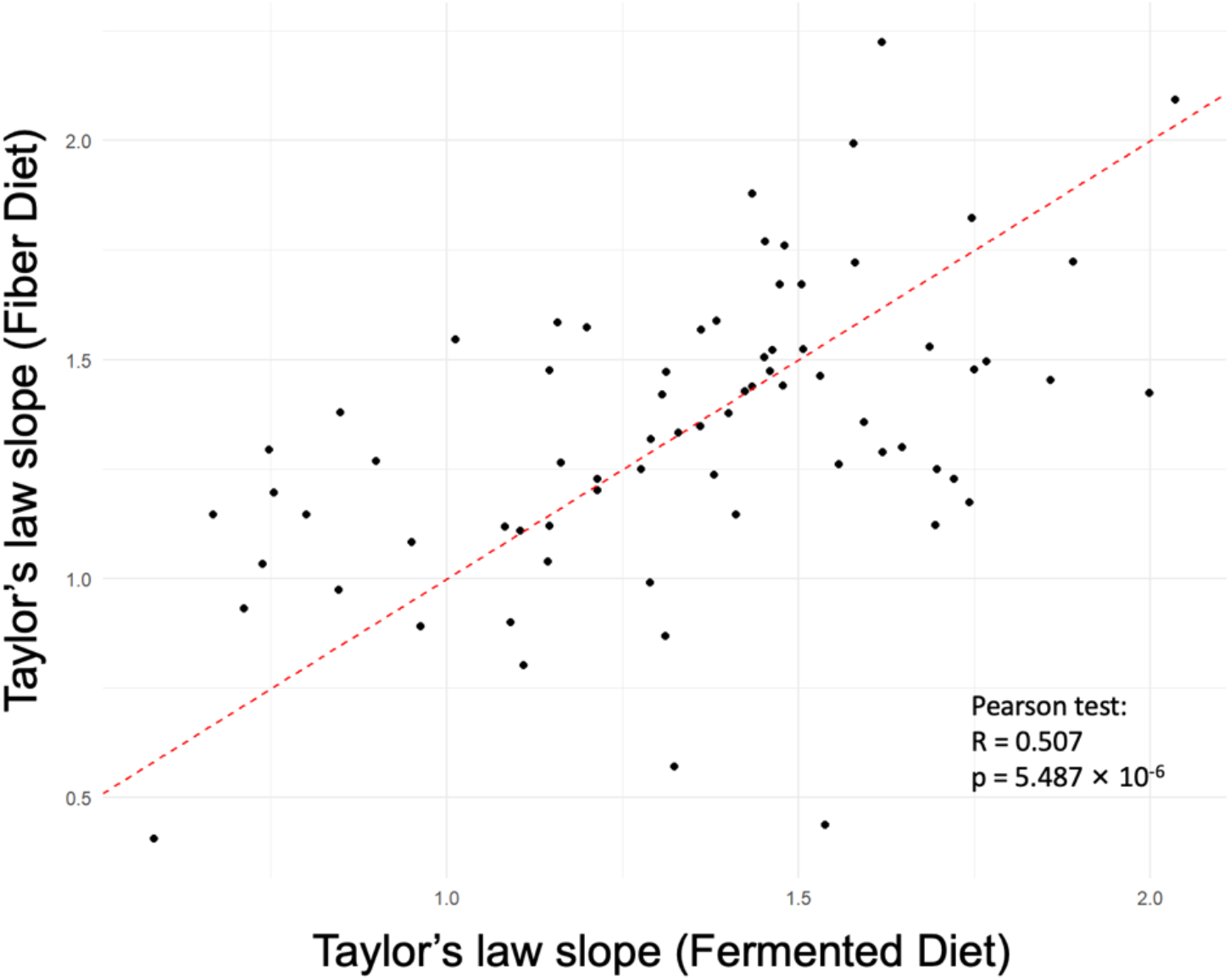
The relationship between species Taylor’s law slopes for each ASV when calculated only using the hosts on either the Fermented or the Fiber diet. Each point represents an ASV, and the dotted line represents the identity line.

**Figure S7.**
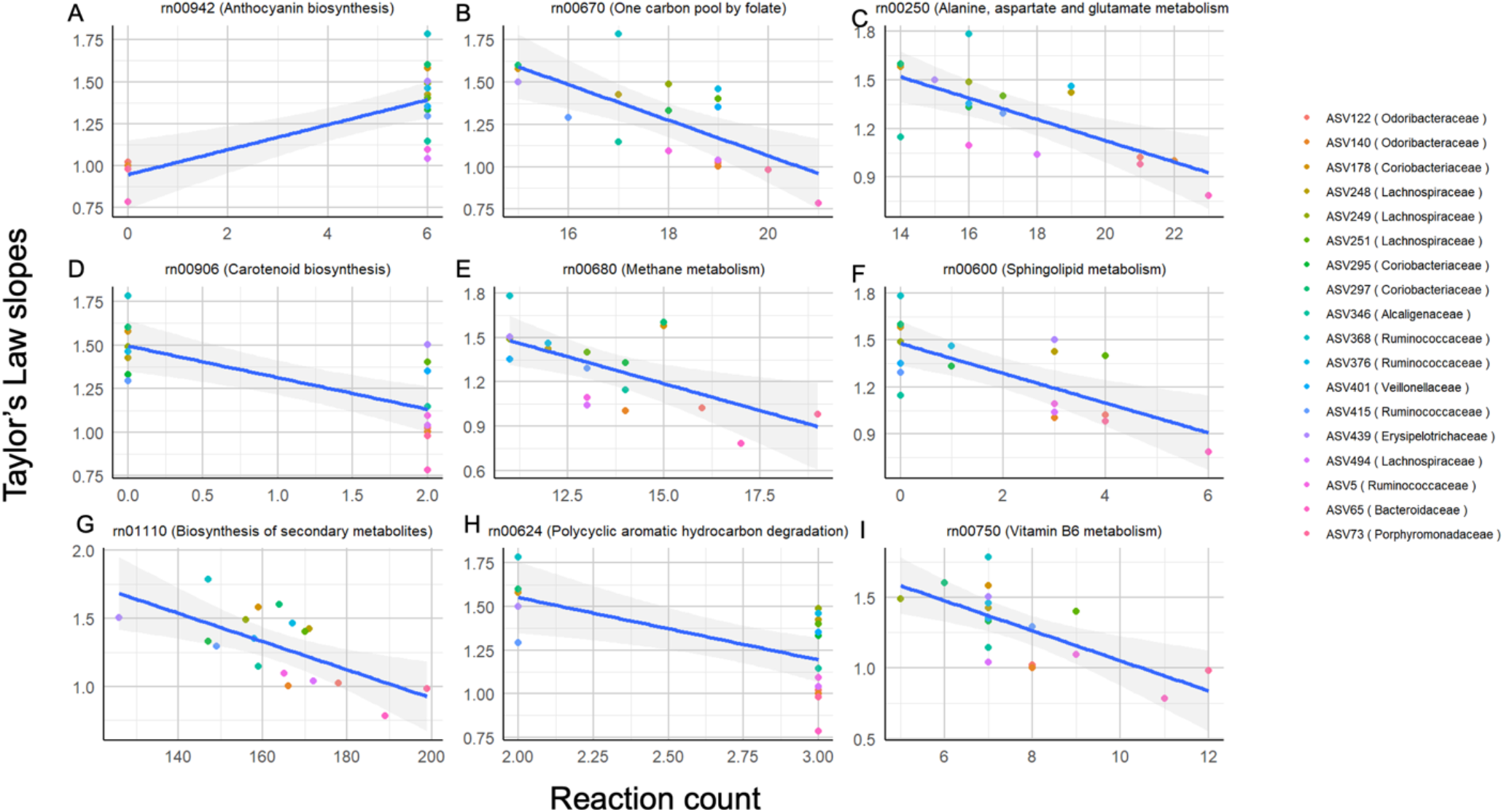
Reaction counts for each KEGG pathway. Each panel shows the relationship between the number of ASV reactions that mapped to each pathway (x-axis) with the species Taylor’s law slope of that ASV (y-axis), across ASVs (points). Data is shown for the 9 KEGG pathways for which this relationship is significant (Pearson correlation test p< 0.05).

